# Exosome based multivalent vaccine: achieving potent immunization, broadened reactivity, and strong T cell responses with nanograms of proteins

**DOI:** 10.1101/2023.01.10.523356

**Authors:** Mafalda Cacciottolo, Justin B Nice, Yujia Li, Michael J. LeClaire, Ryan Twaddle, Ciana Mora, Stephanie Y. Adachi, Esther R. Chin, Meredith Young, Jenna Angeles, Kristi Elliott, Minghao Sun

**Affiliations:** Capricor Therapeutics, Inc., 10865 Road to the Cure, San Diego, CA, 92121

**Author notes:** Corresponding author: Minghao Sun, PhD, Vice President, Research & Product Development, Capricor Therapeutics, Inc. 10865 Road to Cure, suite 150, San Diego, CA 92121. M.C and J.B.N contributed equally to this work.

**Keywords:** exosome, SARS-CoV-2, severe respiratory syndrome coronavirus 2, spike, nucleocapsid, neutralizing antibodies, omicron, lentiviral system, COVID, vaccine, therapeutic

## Abstract

Current approved vaccines against severe acute respiratory syndrome coronavirus 2 (SARS-CoV-2) have focused solely on the spike protein to provide immunity. The first vaccines were developed rapidly using spike mRNA delivered by lipid nanoparticles but required ultra-low storage and have had limited immunity against variations in spike. Subsequently, protein-based vaccines were developed which offer broader immunity but require significant time for development and use of an adjuvant to boost immune response. Here, exosomes were used to deliver a bi-valent protein-based vaccine, in which two independent viral proteins were used. Exosomes were engineered to express either SARS-CoV-2 Delta spike (Stealth X-Spike, STX-S) or the more conserved nucleocapsid (Stealth X-Nucleocapsid, STX-N) protein on the surface. When administered as single product (STX-S or STX-N) or in combination (STX-S+N), both STX-S and STX-N induced a strong immunization with the production of a potent humoral and cellular immune response. Interestingly, these results were obtained with administration of only nanograms of protein and without adjuvant. In two independent animal models (mouse and rabbit), administration of nanograms of the STX-S+N vaccine resulted in increased antibody production, potent neutralizing antibodies with cross-reactivity to other variants of spike and strong T-cell responses. Importantly, no competition in immune response was observed, allowing for delivery of nucleocapsid with spike to offer improved SARS-CoV-2 immunity. These data show that the StealthX^TM^ exosome platform has an enormous potential to revolutionize vaccinology by combining the advantages of mRNA and recombinant protein vaccines into a superior, rapidly generated, low dose vaccine resulting in potent, broader immunity.

## INTRODUCTION

The pandemic emergency has brought to light the need for better vaccines, which can be rapidly generated to protect the population broadly against viruses and their consequent morbidity. More recently, increasing cases of severe acute respiratory syndrome coronavirus 2 (SARS-CoV-2) caused by constantly changing variants of spike, together with the surge of influenza and respiratory syncytial viruses (RSV) has demonstrated the urgent need for multi-valent vaccines which could improve SARS-CoV-2 immunity while also preventing other viruses.

Exosomes offer a new antigen delivery system which could be utilized to rapidly generate multi-valent protein-based vaccines. Exosomes, first identified as extracellular membrane vesicles [1, 2], are small vesicles of ~30-200 nm diameter, enriched in specific subsets of proteins (CD81, CD63, CD9; Tetraspanin family), RNAs, and lipids [3] and responsible for cell to cell communication. Being an endogenous cell product, exosomes have extremely low immunogenicity and toxicity. Purification and scale-up processes for exosomes rely on well-established particle protocols utilizing bioreactors and differential filtration for the manufacturing process. Thus, exosomes can be rapidly engineered to express targets of interest and easily produced for use in multi-valent vaccines.

SARS-CoV-2 has been the promoter of scientific advancement in the vaccinology field, resulting in the rapid development and approval of new mRNA-based vaccines from Moderna and Pfizer [4, 5] and subsequent approval of protein-based vaccines from NovaVax [6, 7]. These vaccines were developed to direct the immune response to the surface spike protein, which binds to the host cell receptor angiotensin-converting enzyme 2 (ACE2), mediating viral cell entry and infection [8]. The spike (S) protein, a class I fusion glycoprotein and major surface protein on the SARS-CoV-2 virus, mediates the binding to the ACE receptor on cell surface promoting its propagation and infection, and it is the primary target for neutralizing antibodies. Since the beginning of the COVID-19 pandemic, the SARS-CoV-2 virus has mutated extensively, reducing the efficacy of current vaccines, in particular mRNA-based vaccines, and creating the need for more effective approaches.

The SARS-CoV-2 genome encodes for three additional structural proteins, important for the virus biology: nucleocapsid, envelope and membrane. Nucleocapsid (N, Ncap) protein, contrary to spike, is highly conserved, with 90% amino acid homology and fewer mutations over time [9]. Ncap binds to the mRNA during the packaging of the RNA genome, regulates viral RNA synthesis during replication and transcription, and modulates its metabolism in infected subjects. IgG antibodies against Ncap have been detected in SARS patients [10, 11], with a significant prognostic value [11]. Despite not being a target of neutralizing antibodies, Ncap protein can stimulate the T-cell response [12] and improve the efficacy of next generation vaccines [13].

While the mRNA-based vaccines were of undeniable help during the emergency, they lack long-term protection [14], likely due to unknown translation efficiencies and the lack of cross-reactivity against new variants of concern (VOC). Thus, there is a pressing demand to develop next-generation COVID-19 vaccine strategies [15]. While many approaches could be taken in consideration, updating the spike antigen to match the circulating VOC seems to be the easiest option: this may result in inaccurate identification of the right variant and therefore lack of vaccine efficacy, as observed for influenza virus and more recently SARS-CoV-2 [16]. An alternative approach could reside in a “mix and match” approach, for which the spike antigen is formulated together with a more conserved viral antigen, such as nucleocapsid for SARS-CoV-2. This approach would broaden the immune response at both the humoral and T cell level, and result in more efficient protection against current and future forms of the virus.

We have previously shown that immunization with our exosome-based SARS-CoV-2 Delta spike vaccine was able to induce a potent, broad immune response (Cacciottolo et al, under review). These results were achieved with only nanograms of spike protein delivered by exosomes without any adjuvants. Therefore, by utilizing our exosome platform (StealthX™, STX) to engineer exosomes to express SARS-CoV-2 Delta spike protein (STX-S) or nucleocapsid protein (STX-N) on their surface, we were able to produce a cocktail of exosomes to create a bi-valent vaccine against SARS-CoV-2, named STX-S+N. Administration of the STX-S+N vaccine in mice and rabbits resulted in an increase of both CD4^+^ and CD8^+^ T cell responses, accompanied by a potent humoral immune response as demonstrated by high levels of neutralizing antibodies, not only against the delta SARS-CoV-2 virus but also the Omicron variants. Importantly, these results were observed after STX vaccination using only nanogram amounts of SARS-CoV-2 proteins presented by exosomes in absence of any adjuvant or lipid nanoparticles, which further strengthens the STX exosome safety profile as a vaccine candidate. Additionally, due to the significantly lower amount of protein (nanograms for STX vs milligrams for standard recombinant protein vaccines) required to cause a robust immune response, this study opens the door to multivalent vaccines using multiple SARS-CoV-2 antigens in combination with influenza and/or RSV for a much broader and safer future immunization campaign.

These results support the clinical development of the STX vaccine for immunization against COVID-19.

## METHODS

### Cell lines

Human embryonic kidney 293 T cells (293T) were purchased from ATCC (CRL-3216). 293T cells were maintained in culture using Dulbecco’s Modified Eagle Medium (DMEM), high glucose, Glutamax™ containing 10% fetal bovine serum. 293T cells were incubated at 37°C /5% CO_2_. FreeStyle™ 293F cells (Gibco, 51-0029) were purchased from ThermoFisher. 293F cells were used as a parental cell line to generate spike SARS-CoV-2 delta spike and nucleocapsid expressing stable cell lines: Stealth X-Spike (STX-S) and StealthX-Nucleocapsid (STX-N) cells. Parental and engineered 293F cells were maintained in a Multitron incubator (Infors HT) at 37°C, 80% humidified atmosphere with 8% CO_2_.

### Lentiviral vectors

Lentiviral vectors for expression of SARS-CoV-2 spike (Delta variant B.1.617.2, NCBI accession # OX014251.1) and Sars-CoV-2 nucleocapsid (NCBI accession # OP359729.1) were designed and synthesized from Genscript together with the two packaging plasmids (pMD2.G and psPAX2). Lentiviral particles for transduction were generated as previously described (Cacciottolo et al, under revision). Briefly, lentiviral particles for transduction were generated by transfecting 293T cells with pMG.2 (Genescript), psPAX2 (Genescript) and STX-S_pLenti (Genscript) expressing spike or STX-N_pLenti (Genscript) expressing nucleocapsid using Lipofectamine 3000 according to the manufacture’s instruction. Spike and nucleocapsid lentiviral particles were collected at 72 hours post transfection and used to transduce 293F parental cells to generate STX-S and STX-N respectively.

### Flow cytometry

Standard flow cytometry methods were applied to measure the spike SARS-CoV-2 protein expression on STX cell surface. In brief, 250K STX cells were aliquoted, pelleted and resuspended in 100uL eBioscience™ Flow Cytometry Staining Buffer (ThermoFisher). Cells were incubated at room temperature (RT) for 30 min protected from light in the presence of anti-spike antibody (Abcam, clone 1A9, ab273433) or anti-nucleocapsid (Abcam, ab281300) labeled with Alexa Fluor®-647 (Alexa Fluor® 647 Conjugation Kit (Fast)-Lightning-Link® (Abcam, ab269823) according to the manufacturer’s protocol. Following incubation, STX cells were washed with eBioscience™ Flow Cytometry Staining Buffer (ThermoFisher, cat No 00-4222-57), resuspended in PBS and analyzed on the CytoFlex S flow cytometer (Beckman Coulter). Data was analyzed by FlowJo (Becton, Dickinson and Company; 2021).

### Cell sorting

Cell sorting was performed at the Flow Cytometry Facility at the Scripps Research Institute (San Diego, CA). To enrich the spike positive population, STX-S cells were stained as described above for flow cytometry and went through cell sorting (Beckman Coulter MoFlo Astrios EQ) to generate pooled STX-S. The pooled STX-S were used in all the experiments in this paper unless specified otherwise.

### STX exosome production

STX-S and STX-N cells were cultured in FreeStyle media (ThermoFisher, 12338018) in a Multitron incubator (Infors HT). Subsequently, cells and cell debris were removed by centrifugation, while microvesicles and other extracellular vesicles larger than ~220 nm were removed by vacuum filtration. Next, exosomes were isolated using either Capricor’s lab scale or large-scale purification method. For lab scale: supernatant was subjected to concentrating filtration against a Centricon Plus-70 Centrifugal filter unit (Millipore, UFC710008), then subjected to size exclusion chromatography (SEC) using a qEV original SEC column (Izon, SP5). For large scale: supernatant was subjected to concentrating tangential flow filtration (TFF) on an AKTA Flux s instrument (Cytiva, United States) and then subjected to chromatography on an AKTA Avant 25 (Cytiva, United States).

### Nanoparticle tracking analysis

Exosome size distribution and concentration were determined using ZetaView Nanoparticle Tracking Analysis (Particle Metrix, Germany) according to manufacturer instructions. Exosome samples were diluted in 0.1 μm filtered 1X PBS (Gibco, 10010072) to fall within the instrument’s optimal operating range.

### Jess automated western blot

Detection of SARS-CoV-2 spike and nucleocapsid proteins in cell lysate and exosomes used Protein Simple’s Jess capillary protein detection system. Samples were lysed in RIPA buffer (ThermoFisher Scientific, 8990) supplemented with protease/phosphatase inhibitor (ThermoFisher Scientific, A32961), quantified using the BCA assay (ThermoFisher Scientific, 23227) and run for detection. To detect spike, the separation module 12–230 kDa was used following manufacturers protocol. Briefly, 0.8 μg of sample and protein standard were run in each capillary, probed with anti-mouse Ms-RD-SARS-COV-2 (R&D Systems, MAB105401, 1:10 dilution) or anti-rabbit Nucleocapsid (NovusBiologics, NBP3-00510, 1:100 dilution) followed by secondary antibody provided in Jess kits (HRP substrate used neat).

### TEM imaging for characterization of STX exosome morphology

STX-S and STX-N exosome samples were negatively stained onto copper grids with a carbon film coating and imaged by TEM at the Electron Microscopy Core Facility at UC San Diego (San Diego, CA). Briefly, samples were treated with glow discharge, stained with 2% uranyl acetate and dried before imaging. Grids were imaged on a JEM-1400 Plus (JEOL Ltd, Japan) at 80kV and 48uA. Images were taken at from 12K-80K magnification at 4kx4k pixels of resolution.

### CD81 bead-assay

STX-S or 293F parental exosomes were mixed with anti-CD81 labeled magnetic beads for 2 h at RT (ThermoFisher, 10622D) and washed twice with PBS using the magnetic stand. Next, the bead-exosomes were incubated with either direct conjugated AlexaFluor 647 anti-spike (see above flow cytometry), FITC anti-CD81 antibody (BD Biosciences, 551108), or FITC Mouse IgG, κ isotype control (BD Biosciences, 555748) for 1 h at RT followed by two PBS washes. 293F exosomes were used as a negative control for spike expression and the isotype antibody was used as a negative control for CD81 expression. Samples were analyzed on a CytoFlex S (Beckman Coulter) flow cytometer and data was analyzed by FlowJo.

### Animal studies - Mice

To examine the efficacy of STX exosomes, age matched BALB/c mice (female, 8-10 wks old) were anesthetized using isoflurane and received bilateral intramuscular injection (50 μl per leg, total 100 μl) of either 1) PBS 2) STX-S, 3) STX-N or 4) STX-S+N exosomes. Booster injection was performed at day 21. Mice were monitored closely for changes in health and weight was recorded biweekly. Blood collection was performed at day 14 and day 35. Blood (~50-500 μl) was collected from the submandibular vein and processed for plasma isolation after centrifugation at 4000 rpm for 5 min at 4°C. For comparison study, mice were injected with equal amounts of SARS-CoV-2 protein delivered as 1) soluble protein in conjugation with adjuvant (Alhydrogel, 100 μg/dose, vac-alu-250, InvivoGen), or 2) STX exosome. Blood was collected 2 weeks after injection and tested for IgG against SARS-CoV-2. Timeline of mouse study is outlined in Fig. 3A. Mouse tissues (brain, salivary gland, heat, lung, liver, spleen, kidney, gastro-intestinal tract (GI) and skeletal muscle (site of injection) were collected by the Capricor Therapeutic’s team and fixed in fixed in 10% Neutralized Formalin. Tissues were sent to Reveal Biosciences for further processing and analysis. Sections were stained with hematoxylin and eosin and analyzed for alterations. Pathology evaluation was performed by Dr Mary E.P. Goad (DVM, PhD, DACVP, DACLAM, DABT).

### Animal studies – Rabbits

Study was conducted at CBSET, Inc. (CBSET), accredited by AAALAC International. All procedures and conditions of testing followed the USDA and AWA1/AWR2 guidelines. To evaluate the potential toxicity and host immune response of STX-S+N vaccine, age matched rabbits (male/female, New Zealand White, 2.5-3.0 kg) received intramuscular (IM) injection of the intended human dose (10-200 ng total protein in 0.5 mL) of the STX-S+N vaccine. Control animals received phosphate-buffered saline (PBS). Booster injection was performed at day 14. Rabbits were monitored closely for changes in health and weight recorded. Blood collection was performed weekly on day 0, 7, 14, 21, 28 and processed for plasma isolation after centrifugation at 4000 rpm for 5 min at 4°C. Timeline of rabbit study is outlined in Fig. 4A. Rabbit tissues were collected by the CBSET team at terminal (day 28), fixed in 10% Neutralized Formalin and processed for pathological alterations.

### IgG ELISA

Mouse and Rabbit IgG antibody against SARS-CoV-2 spike or nucleocapsid was measured by enzyme-linked immunosorbent assays (ELISA) using precoated ELISA plates (IEQ-CoV-S-RBD-IgG and IEQ-CoV-N-IgG, RayBiotech) according to the manufacturer’s instructions, at RT. Briefly, plasma samples were diluted in sample buffer (RayBiotech) and added to antigen-coated wells, in triplicates, and incubated at RT for 2 h on a shaker (200 rpm). Commercially available antibody against Spike (S1N-S58, Acro Biosystems) or Nucleocapsid (NUN-S47, Acro Biosystems) was used as positive controls. Plates were washed 3 times with wash buffer, incubated for 1h at RT with HRP-conjugated goat anti-mouse secondary antibodies (115-035-003, dilution 1:5000, Jackson ImmunoResearch) or anti-rabbit (111-035-003, dilution 1:5000, Jackson ImmunoResearch) diluted in assay buffer (RayBiotech). After 3 washes, plates were developed using TMB substrate (RayBiotech). After 15 min incubation, reaction was stopped by adding STOP solution and absorbance at 450 nm was recorded using a BioTeck Gen5 plate reader (Agilent). Endpoint titers were calculated as the dilution that emitted an optical density exceeding 4X the PBS control group.

### Neutralizing antibodies against DELTA SARS-CoV-2

Samples were analyzed by Retrovirox, Inc. (San Diego, CA). Briefly, Vero E6 cells were used to evaluate the neutralization activity of the test-items against a replication competent SARS-CoV-2 delta variant (B.1.617.2). Samples were pre-incubated with the virus for 1 h at 37°C before addition to cells. Following pre-incubation of plasma/virus samples, cells were challenged with the mixture. Samples were present in the cell culture for the duration of the infection (96 h), at which time a *Neutral Red* uptake assay was performed to determine the extent of the virus-induced cytopathic effect (CPE). Prevention of the virus-induced CPE was used as a surrogate marker to determine the neutralization activity of the test-items against SARS-CoV-2. Test-items were evaluated in duplicates using two-fold serial dilutions starting at a 1:40 dilution (eight total dilutions). Control wells included “CoV02-Delta” and GS-441524, tested in singlet data points on each plate. “CoV02-Delta” is convalescent plasma from an individual who previously received two doses of the Moderna COVID-19 vaccine before infection with the Delta variant. GS-441524 is an antiviral from Gilead Sciences. CoV-01 is plasma from a patient who received two doses of the Moderna covid-19 vaccine with no prior history of SARS-CoV-2 infection. NT50 values of the test-items were determined using *GraphPad Prism* software. Data are available in Final_Report_CAPR_2022_04.

### Neutralizing antibodies against Omicron (BA.1 and BA.5.2.1) SARS-CoV-2

Samples were analyzed by Retrovirox, Inc. (San Diego, CA). Neutralization assays were performed using anti-NP immunostaining (Omicron BA.1 and BA.5.2.1). Briefly, samples were pre-incubated with the virus for 1 h at 37°C before addition to Vero E6 cells. Following incubation, media was removed, and then cells were challenged with the SARS-CoV-2 / test-item pre-incubated mix. The amount of viral inoculum was previously titrated to result in a linear response inhibited by antivirals with known activity against SARS-CoV-2. Cell culture media with the virus inoculum was not removed after virus adsorption, and test-items and virus were maintained in the media for the duration of the assay (48 h). Subsequently, the extent of infection is monitored by incubating cells with a monoclonal test-item against the SARS-CoV-2 nucleocapsid (NP). The amounts of the viral antigen in infected cells are estimated after incubation with horseradish peroxidase conjugated polyclonal test-items against human IgG (HRP-goat anti-mouse IgG). The reaction is monitored using a colorimetric readout (absorbance at 492nm). Test-items were evaluated in duplicates using two-fold serial dilutions starting at a 1:40 dilution. Control wells included GS-441524 (Gilead Sciences), tested in singlet data points on each plate. Data are available in Final_Report_CAPR_2022_04.

### Splenocyte isolation

Spleens were processed for single cell isolation by mechanical disruption of spleen pouch using a syringe stopper and passage through a 0.040 mm mesh size nylon cell strainer to remove tissue debris. Erythrocytes were lysed using ammonium chloride potassium (ACK) buffer (A1049201, ThermoFisher), splenocytes were collected by centrifugation. Cellular pellet was resuspended in complete RPMI 1640 media (FG1215, Millipore Sigma Aldrich).

### ELISPOT

Briefly, splenocytes were isolated by mechanical disruption of spleen pouch and seeded at a concentration of 5E5 cells/well and incubated for 24 h in the presence or absence of 10 μg/ml of SARS-CoV-2 Spike (S1N-C52H4, AcroBiosystems) or Nucleocapsid (NUN-C5227, AcroBiosystems). Commercially available ELISPOT plates for evaluation of IL-4 (MuIL4, Immunospot, Cellular Technology Limited) and IFNg (MuIFNg, Immunospot, Cellular Technology Limited; cat# 3110-4APW-10, rabbit, MabTech) were used. Assay was performed according to manufacturer’s guidelines. Plates were analyzed using the ELISPOT reader S6ENTRY (Immunospot, Cellular Technology Limited).

### ELISA for protein quantification

Spike protein level on exosomes was measured by ELISA using precoated ELISA plates (ELV-COVID19S1, RayBiotech) according to the manufacturer’s instructions, at RT. Nucleocapsid level on exosomes was measured by ELISA using precoated ELISA plates (Legend Max SARS-CoV2 Nucleocapsid protein ELISA Kit, 448007, BioLegend) according to the manufacturer’s instructions, at RT. Briefly, samples and standards were loaded to the precoated plate, and incubated 2-2.5 hr at RT on a shaker (200 rpm). Plates were washed and incubated with biotin-conjugated detection antibody for an hour at RT, followed by 45 min incubation in streptavidin solution. After washes, plates were developed using TMB substrate. After 30 min incubation, reaction was stopped by adding STOP solution and absorbance at 450 nm was recorded using a BioTeck Gen5 plate reader (Agilent). For nucleocapsid, after 10 min incubation in TMB substrate, absorbance was recorded at 450 nm and 570nm. For analysis, the absorbance at 570 nm can be subtracted from the absorbance at 450 nm, and OD used to build a standard curve.

### Pathology

Mouse tissues (brain, salivary gland, heat, lung, liver, spleen, kidney, gastro-intestinal tract (GI) and skeletal muscle (site of injection) were collected by the Capricor Therapeutic’s team and fixed in fixed in 10% Neutralized Formalin. Tissues were sent to Reveal Biosciences for further processing and analysis. Sections were stained with hematoxylin and eosin and analyzed for alterations. Pathology evaluation was performed by Dr Mary E.P. Goad (DVM, PhD, DACVP, DACLAM, DABT).

### Statistical analysis

Data were analyzed using Excel and GraphPad Prism 9.1 and shown as mean±sem. 1-way ANOVA with post-hoc correction for multiple comparisons or 2-tailed t-test were applied as needed.

## RESULTS

### SARS-CoV-2 protein expression on the surface of STX producing cells and exosomes

STX cells were generated by lentiviral transduction and expression of SARS-CoV-2 proteins on cells surface was evaluated by flow cytometry (Figure 1). As shown, STX cells showed > 95% increased expression of spike and nucleocapsid (Fig. 1A) compared to parental 293F cells.

**Figure 1.**
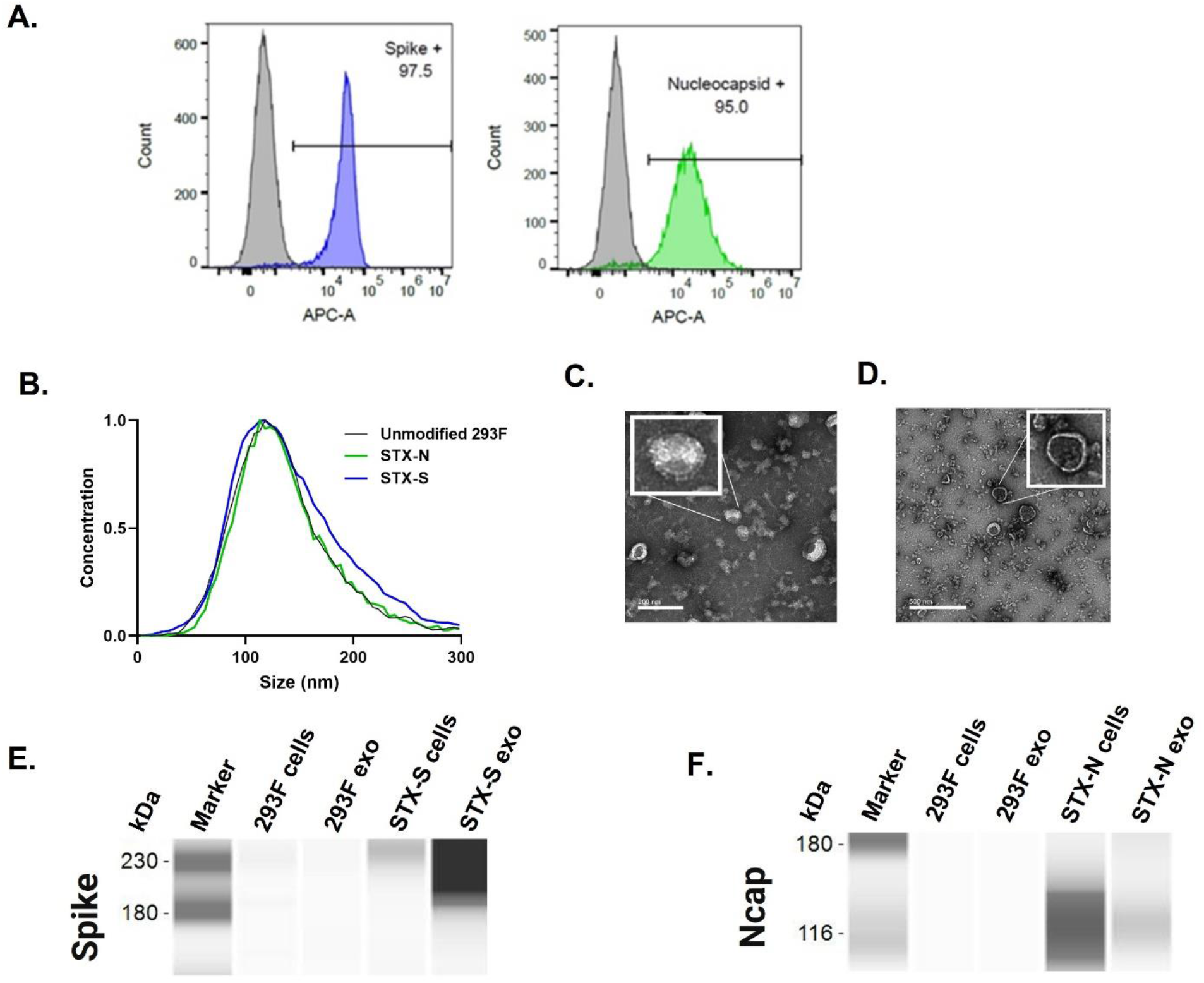
Characterization of SARS-CoV-2 proteins on cell and exosome surface. **A.** Representative images of gating strategy for protein expression analysis on cell membrane. High expression of spike (**blue histogram**) and nucleocapsid (**green histogram**) was detected on cell surface by flow cytometry. Parent, non-engineered 293F cells do not express SARS-CoV-2 antigens, as expected (**grey histogram**). **B.** Size distribution of STX-S and STX-N exosomes by ZetaView nanoparticle tracking analysis (NTA). **C-D.** TEM image of purified STX-S (**C**) and STX-N (**D)** exosomes. **E-F.** Spike **(E)** and nucleocapsid **(F)** expression analyzed by Jess-automated Western Blot. 0.8ug protein was loaded per lane, as calculated by BCA assay.

STX exosomes were purified from the engineered 293F cell culture supernatant using Capricor’s lab-scale purification technique as previously reported (Cacciottolo et al, under review; see methods).

The purified STX-S and STX-N exosomes showed an expected average diameter of 144.6 nm and 140.4 nm (Fig. 1B) and an expected polydispersity index (PDI) of <0.2 (0.152 and 0.129, Fig. 1B), respectively [17].

STX exosomes were analyzed by TEM imaging. As shown in Fig. 1B-C, typical exosome size and morphology were observed with round, smooth nano particles detected that had a visible lipid bilayer. Importantly, spike protrusions were visible on the surface of the STX-S nanoparticles indicating the presence of spike protein on exosomes (Fig. 1C). For Nucleocapsid, a characteristic lipid bilayer was observed, which resulted thicker than naïve exosomes, suggesting an accumulation of particles in the exosome membrane (Fig. 1D).

Spike and Nucleocapsid expression were validated on cell lysates and exosomes using Protein Simple’s Jess automated Western blot. Spike protein was detected in both STX-S cells and exosomes, with enrichment of spike protein in the exosome samples. STX-N engineered cells and exosomes expressed abundant levels of Nucleocapsid (Fig. 1E-F). Further, SARS-CoV-2 protein spike and Ncap were detected on exosome membrane using a bead-based CD81 assay, with more >75% expression together with exosome specific marker CD81.

The concentration of Spike antigen in STX-S exosomes and Nucleocapsid antigen in STX-N exosome was further quantified by ELISA. A final STX-S prep of 1×10^12 exosomes/mL contains on average 253.77 ng of Spike, while a final STX-N prep of 1×10^12 exosomes/mL contains on average 40.59 ng of nucleocapsid.

### STX-S and STN-N individually induced strong immunization, in absence of adjuvant

To validate the capability of STX-S and STX-N exosomes to induce an immune-response, spike and nucleocapsid, mice were immunized with 10ng exosome formulation STX-S and STX-N. As comparison to show the robustness of exosome delivery, 10 ng of either spike or Ncap recombinant protein delivered in conjugation with adjuvant (Alhydrogel, InvivoGen). PBS was used as control. Blood collected 2 weeks after the boost injection (2^nd^ injection) showed that both STX-S and STX-N vaccines increased antibody production respectively against spike and nucleocapsid in all animals, as expected based on previous studies (Cacciottolo et al, under review). Neither spike or Ncap protein in combination with adjuvant were statistically different than the PBS control (negative control), and no antibody production was observed (Fig. 2).

**Figure 2.**
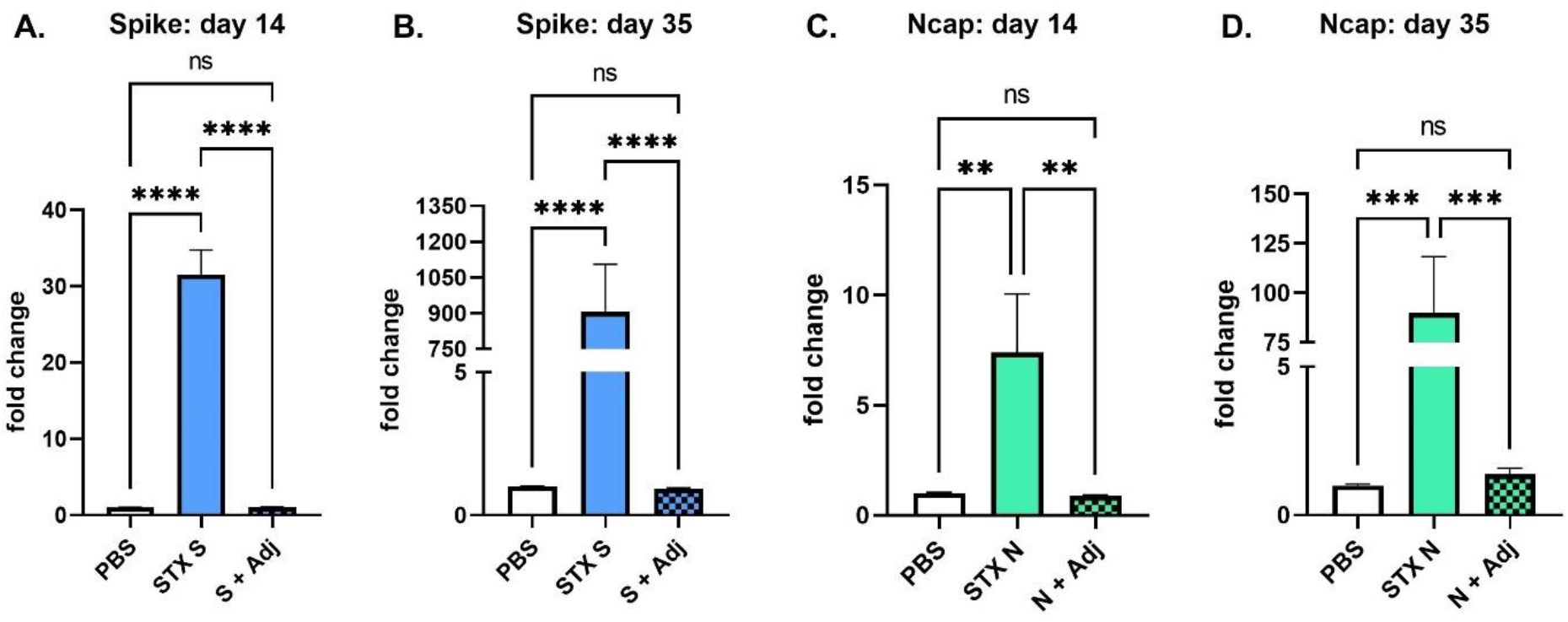
STX-S and STX-N exosome induce strong immunization without adjuvant. **A.** Increased IgG production against spike after 1 i.m. injection. **B.** Increased IgG production against spike after 2 i.m. injection. **C.** Increased IgG production against Ncap after 1 i.m. injection. **D.** Increased IgG production against Ncap after 2 i.m. injection. Spike (S + Adj) and Ncap (N+Adj) protein at the same dose in combination with adjuvant. N= 10/experimental group. Data are shown as mean ± SEM. **** p<0.0005, ***p<0.001, **p<0.01, ns= not significant, 1-way ANOVA

### STX-S+N vaccine induces strong immunization against SARS-CoV2 protein in mice

We proceeded to assess the immune response of a multivalent vaccine, obtained by the combination of STX-S and STX-N exosomes. STX-S+N was administered at three different doses (Table 1) into mice by two i.m. injections. A second i.m. injection, boost injection, was delivered after a 3-week interval. PBS was used as negative controls in the study.

**Table 1.** Concentration of Spike and Ncap used in the study.

|  | Spike ng/injection | Ncap ng/injection |
| --- | --- | --- |
| **Dose 1** | 25 | 2.5 |
| **Dose 2** | 10 | 4 |
| **Dose 3** | 3 | 9 |

Immunization to the STX-S+N vaccine was evaluated by the quantification of antibodies against Ncap and Spike (Fig. 3B-E). Increase of antibody production was detected after the first injection and continued to increase after the boost injection (2^nd^ injection). A single injection of STX-S+N induced an increase up to 30-fold in IgG against spike, with overall no significant difference between the three dose. After complete immunization, dose 1 (25ng/spike) and dose 2 (10ng/spike) resulted in a 1500-fold increase in antibody against Spike, while dose 3 (3ng/spike, significantly lower amount of spike) resulted in a 280-fold increase. On the other hand, a dose response was observed for Ncap. An increase of 1.5-fold was observed for the low dose 1 (2.5ng/Ncap), ~4-fold increase for dose 2 (4ng/Ncap) and up to 7-fold for dose 3 (9ng/Ncap). After complete immunization cycle (2 i.m. injections) no significant difference was observed across doses, with an increase in IgG against Ncap between 24-43-fold increase observed in STX-S+N treated mice.

**Figure 3.**
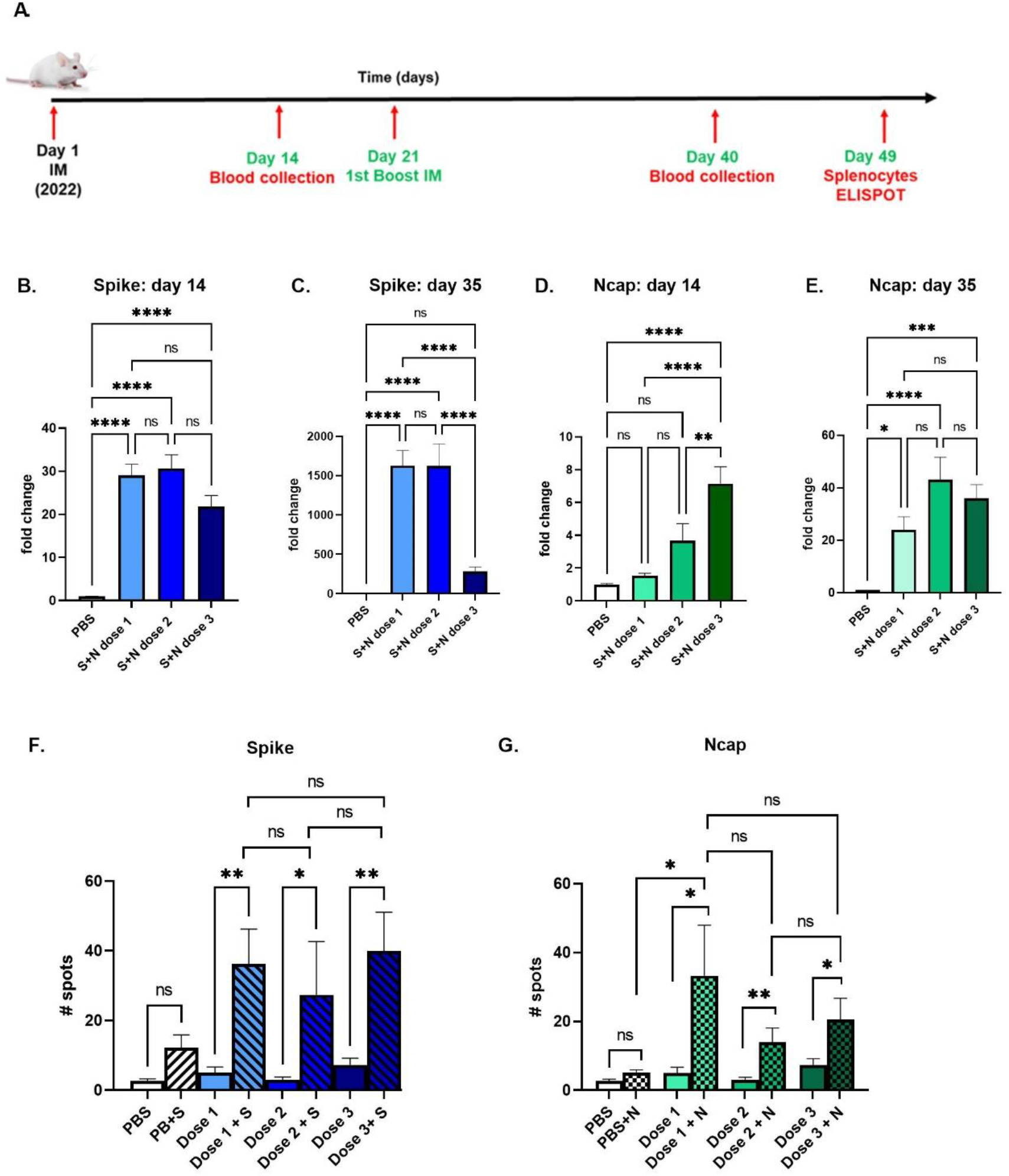
STX-S+N vaccine elicits a strong antibody and cellular response in mice. STX-S+N vaccine induced increase of antibodies against SARS-CoV-2 Spike and Nucleocapsid, and a strong T-cell response. **A.** Timeline of mouse study. **B.** IgG against Spike at day 14. **C.** IgG against Spike at day 35. **D.** IgG against Ncap at day 14. **E.** IgG against Ncap at day 35. PBS was used as a vehicle control. **F.** IFNγ response is increased after Spike stimulation. **G.** INFγ response is increased after Ncap stimulation. Data are shown as mean ± SEM. *p<0.05, **p<0.01, ***p<0.005, **** p<0.001, ns= not significant, 1-way ANOVA, adjusted for multiple comparison. N= 10 animals per experimental group.

To characterize the T cell response to STX-S+N, antigen-specific T cell responses were measured by ELISpot (Fig. 3F-G). Vaccination with STX-S+N elicited multi-functional, antigen-specific T cell responses. Splenocytes were isolated from animals at day 35 (2 weeks after boost (2^nd^) injection) and evaluated using ELISpot plates precoated with IFNγ. PBS was used as controls in the study as described above. Baseline expression was compared to stimulation with 10 μg/ml of either Spike or Ncap protein (AcroBiosystem). While baseline IFNγ response was comparable between groups, evaluation of IFNγ secreting cells in response to ex vivo of either Spike (Fig. 3F) or Ncap (Fig. 3G) stimulation showed a strong increase in spleens immunized with STX-S+N vaccine, suggesting a Th1-biased CD8+ T cell response. After spike stimulation, an average of 7-fold increase in IFNγ response was observed despite the STX-S+N dose administered (Fig. 3F). After Ncap stimulation, a dose response effect was observed with 7-fold increase in mice receiving the lowest Ncap dose (dose 1, 2.5ng) and ~3-fold increase in mice receiving either dose 2 (4ng Ncap) or 3 (9ng Ncap) (Fig. 3G).

### STX-S+N vaccine induces strong immunization against SARS-CoV2 protein in rabbits

Immune response to STX-S+N vaccine, using a clinically relevant dose, was evaluated in rabbits. STX-S+N vaccine was administered at two different doses (Table 2) into rabbits by two i.m. injections. A second i.m. injection, boost injection, was delivered after a 2-week interval. PBS was used as controls in the study.

**Table 2.** Concentration of Spike and Ncap used in the study.

|  | **Spike ng/injection** | **Ncap ng/injection** |
| --- | --- | --- |
| **Dose 1** | 125 | 10 |
| **Dose 2** | 50 | 20 |

Increased antibody production was detected as early as one week after the first injection and continued to increase after the boost injection (Fig. 4). After complete immunization cycle (two i.m. injections), up to 3600-fold increase in IgG against Spike (Fig. 4B) and up to 170-fold increase in IgG against Ncap (Fig. 4C) were observed in STX-S+N treated rabbits. No significant differences were observed between doses in antibody production against either antigen.

**Figure 4.**
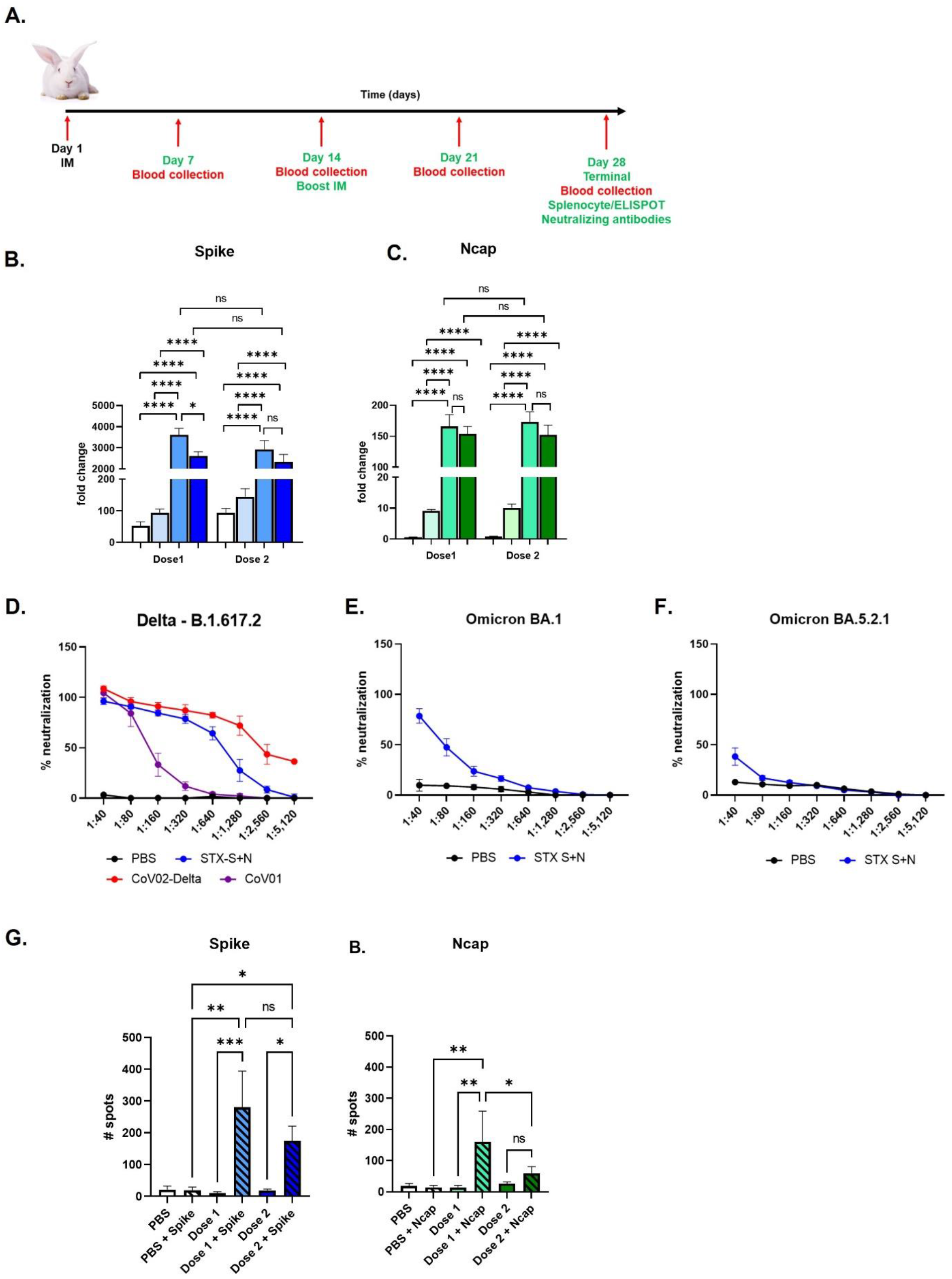
STX-S+N vaccine elicits a strong antibody and T cell response in rabbits. STX-S+N vaccine induced a statistically significant increase of antibodies against SARS-CoV-2 Spike (**B**) and Nucleocapsid (**C**). PBS was used as a vehicle control. Data are reported as fold change over PBS. N= 8 STX-S+N per dose, 8 PBS. **A.** Timeline of Rabbit study. **B.** IgG against Spike **C.** IgG against Ncap. **D-F.** Neutralizing antibodies. **D.** STX-S+N vaccine generated strong neutralization against SARS-CoV-2 delta spike (B.1.617.2) **E.** STX-S+N vaccine resulted in neutralization of SARS-CoV-2 spike Omicron BA. 1 **F.** STX-S+N vaccine resulted in neutralization of SARS-CoV-2 spike Omicron BA.5.2.1. Response after Spike stimulation. **G-H.** T-cell response. **G.** IFNg Response after Spike stimulation. **H.** IFNg Response after Ncap stimulation. Data are shown as mean ± SEM. COV-02-Delta = plasma from a patient immunized with Moderna’s mRNA vaccine with a breakthrough SARS-CoV-2 delta spike infection; CoV-01 = plasma from a patient who received two doses of the Moderna covid-19 vaccine with no prior history of SARS-CoV-2 infection. N= 8 animals per experimental group.

Immunization to the STX-S+N vaccine was further evaluated by assessing neutralizing antibodies against SARS-CoV-2 variants (Fig. 4 D-F). Plasma from rabbits of dose 2 (50ng S and 20ng N, 8 animals) and 2 PBS control were tested for neutralizing antibodies against SARS-CoV-2 Delta variant. Potent neutralizing activity was elicited by STX-S+N in all analyzed animals. Importantly, STX-S+N in rabbits induced a response comparable to the human control plasma (COV-02-delta, plasma from a patient fully immunized with Moderna’s mRNA vaccine, with breakthrough delta infection), with a complete neutralizing response at higher dilutions (i.e., 1:320, range of 116.74 −881.47%; average: 96.22±8.9%) (Fig. 4D). Moreover, STX-S+N in rabbits performed better than CoV-01 control (plasma from a patient who received two doses of the Moderna covid-19 vaccine with no prior history of SARS-CoV-2 infection).

Additionally, the same samples were tested for neutralizing antibodies against SARS-CoV-2 Omicron variants (Omicron BA.1 and BA.5.2.1). As shown in Fig. 4E-F, a strong cross neutralization was observed for the STX-S+N treated rabbits achieving an average neutralization of 75% (range of 40.7-100%) for Omicron BA.1 (Fig. 4E) and a range between 10% to 91% for Omicron BA5 (Fig. 4F). These data suggest that protein-based vaccines delivered by exosomes, specifically STX-S+N, may result in a broader protection against SARS-COV-2 variants. In all assays, rabbits receiving PBS showed no neutralization.

Lastly, T cell response to STX-S+N immunization was measured by ELISpot (Fig. 4 G-H). Vaccination with STX-S+N elicited multi-functional, antigen-specific T cell responses. Splenocytes were isolated from animals at day 28 (2 weeks after boost (2^nd^) injection) and evaluated using ELISpot plates precoated with IFNg. PBS was used as controls in the study as described above. Baseline expression was compared to stimulation with 10 μg/ml of either Spike or Ncap protein (AcroBiosystem). While baseline IFNγ response was comparable between groups, stimulation with either Spike (Fig. 4G) or Ncap (Fig. 4H) resulted in a strong increase in IFNg production in spleens immunized with STX-S+N vaccine, suggesting a Th1-biased CD8+ T cell response. For spike, a clear dose response was observed with greater response in rabbits receiving dose 1 (125ng S, +28-fold) than dose 2 (50ng S, +10-fold), despite not statistically significant (Fig. 4G). For Ncap, dose 1 (10ng N, +10-fold) resulted in better immune response than dose 2 (20ng N, +2-fold) (Fig. 4H).

## DISCUSSION

The pandemic emergency has brought to light the need for a new generation of rapidly developed vaccines that induce a longer lasting, potent, broader immune responses to combat the ever-changing variants of SARS-CoV-2 and hinder virus transmission. While the mRNA vaccines played a critical role during the emergency in reducing SARS-CoV-2 hospitalization rates and deaths, its short-lived immunity and reduced ability to protect against new VOCs, pushes the scientific community to look for new approaches.

A multi-valent, protein-based vaccine delivered by exosomes could meet this urgent need for a new generation of vaccines due to the rapid speed of development, manufacturability, and the ability to produce a strong antibody response, with neutralizing antibodies and a strong T-cell response able to broadly combat viral infection with a minimum number of injections.

Here we have shown that exosomes can be used to deliver viral proteins for immunization: our StealthX™ (STX) platform generated two vaccine candidates (STX-S and STX-N), that independently, and in combination (STX-S+N) induced a strong immune response against two SARS-CoV-2 proteins with a single shot, in two different animal models by delivering nanograms of proteins on the surface of the exosomes. No adjuvant was needed, >100X less protein than traditional recombinant protein vaccines was used, and no competition between proteins was observed.

The “multivalent” or “combination” vaccines have multiple advantages from a public health standpoint [18]. They require fewer shots, which would positively engage the population, increasing the percentage of vaccinated people with broader epidemiological benefits. Consequently, less shots due to combination vaccines also means lower budgeting and cost overall. Of course, there are limitations that need to be taken in consideration. First, selection of antigen: it is crucial that the selected antigens contain as highly conserved sequences as possible with low mutagenesis rates in combination with the most recent, prevalent variants of concern. Importantly, immunogen interference is critical: minimal to no competition should be observed when using a multi-valent vaccine. As per our STX-S+N, the multivalent vaccine shows the same strength of efficacy as the single product, with thousands fold increase in antibody load post-vaccination. Additionally, STX-S+N elicited both a quantitative and qualitative immune response [19]; consistent increase in amount of antibody produced, presence of protection as delineated by neutralizing antibodies and engagement of T cells were all observed in response to the administration of our exosome vaccine STX-S+N. Importantly, no deleterious side effects were recorded: both mice and rabbits showed no changes in weight or blood tests, and no alteration at the tissue levels (data not shown), suggesting an overall safe profile.

Our data suggest that exosomes are ideal vehicles for vaccination because they can safely deliver the antigen of interest (exogenous protein) efficiently by mimicking the natural viral infection. Exosome based vaccines constitute an innovative approach for an efficient virus-free, human-derived vaccine design [19]. Yoo, et. al. observed that exosomes or extracellular vesicles at large could support the vaccination needs beyond traditional strategies: compared to viral or vector methods, exosomes are not immunogenic themselves, but are carrier of a protein that retains the original conformation, tridimensional structure and modifications, all embedded in the lipid biolayer of its membrane and ready to be efficiently presented as such to the immune system [19].

Importantly, the strong T cell response initiated by STX-S+N vaccine administration is clinically relevant and crucial for the new generation of COVID vaccines. Keeton, et. al. observed that while neutralizing antibody might not recognize the new VOC, the T cell population is able to cross react with them and confer protection [20]. We have observed a robust spike- and a nucleocapsid-specific T-cell response: the highly conserved nucleocapsid protein used in the STX-S+N vaccine can further increase and broaden the efficacy of SARS-CoV-2 vaccines, engaging an additional immune response that is not compromised by a surface protein naturally mutating. This suggests that where the new VOC escape the barrier of neutralizing antibodies, the T cell response will likely be cross-reactive and limit the infection [13, 21].

Other groups have reported a multivalent vaccine for SARS-CoV-2 using a viral approach co-expressing the SARS-CoV-2 N and S proteins on VSV virus [22] or by adenovirus [23]. Interestingly, use of a multiprotein vaccine broadens the efficacy to distal organs, reducing the viral charge not only on the respiratory system but to the distant brain, as well [13]. As stated above, nucleocapsid-specific immunity plays an indispensable role during a SARSCoV-2 infection. While antibody responses can block the initial entry of the virus at proximal sites of infection, it is the T cell response that plays a critical role in controlling propagation of infection, second-round infections, and subsequent viral dissemination to distal sites, providing a synergistic antiviral effect by killing virally infected cells and curtailing further the dissemination of the virus to the peripheral organs.

In conclusion, we rapidly generated a single-dose dual-antigen vaccine with efficacy against multiple SARS-CoV-2 VOC, that has broader immune capability, and could be used as a booster vaccine to the existing immunity generated by previously approved vaccines while also delivering nucleocapsid immunity. Our StealthX vaccine technology, which uses rapidly engineered and manufactured exosomes to deliver nanograms of single or multiplexed viral antigens to elicit a strong, broad immune response without any adjuvants or lipid nanoparticles, has the ability to revolutionize the next generation of vaccines.

## Author contribution statements

MC contributed to the design, implementation, and analysis of all research in this paper and to the writing of the manuscript. MS contributed to all the design of STX constructs; JBN, YL produced the cell lines, carried the in vitro experiments, and contributed to the writing of the manuscript. MJL, MY, JA produced the exosomes used in the study. KE, MS contributed to the design and review of studies and the writing of the manuscript. MC, JBN, JL, CM, MJL, JA, MY, SYA, EC, RT carried out the experiments.

The authors declare no conflict of interest.

\# To whom correspondence should be addressed.

## Notes

### Competing Interest Statement

The authors are employees of Capricor Therapeutics.

### Summary of Updates

The manuscript was edited for typos and misspelling errors. No changes were made to the results or conclusions of the paper.

## References

1. Harding, C. and P. Stahl, Transferrin recycling in reticulocytes: pH and iron are important determinants of ligand binding and processing. Biochem Biophys Res Commun, 1983. 113(2): p. 650–8.

2. Pan, B.T. and R.M. Johnstone, Fate of the transferrin receptor during maturation of sheep reticulocytes in vitro: selective externalization of the receptor. Cell, 1983. 33(3): p. 967–78.

3. Gurung, S., et al., The exosome journey: from biogenesis to uptake and intracellular signalling. Cell Commun Signal, 2021. 19(1): p. 47.

4. Baden, L.R., et al., Efficacy and Safety of the mRNA-1273 SARS-CoV-2 Vaccine. N Engl J Med, 2021. 384(5): p. 403–416.

5. Polack, F.P., et al., Safety and Efficacy of the BNT162b2 mRNA Covid-19 Vaccine. N Engl J Med, 2020. 383(27): p. 2603–2615.

6. Vohra-Miller, S. and I.S. Schwartz, NVX-CoV2373, a protein-based vaccine against SARS-CoV-2 infection. CMAJ, 2022. 194(35): p. E1214.

7. Tian, J.H., et al., SARS-CoV-2 spike glycoprotein vaccine candidate NVX-CoV2373 immunogenicity in baboons and protection in mice. Nat Commun, 2021. 12(1): p. 372.

8. Letko, M., A. Marzi, and V. Munster, Functional assessment of cell entry and receptor usage for SARS-CoV-2 and other lineage B betacoronaviruses. Nat Microbiol, 2020. 5(4): p. 562–569.

9. Grifoni, A., et al., A Sequence Homology and Bioinformatic Approach Can Predict Candidate Targets for Immune Responses to SARS-CoV-2. Cell Host Microbe, 2020. 27(4): p. 671–680 e2.

10. Leung, D.T., et al., Antibody response of patients with severe acute respiratory syndrome (SARS) targets the viral nucleocapsid. J Infect Dis, 2004. 190(2): p. 379–86.

11. Batra, M., et al., Role of IgG against N-protein of SARS-CoV2 in COVID19 clinical outcomes. Sci Rep, 2021. 11(1): p. 3455.

12. Okada, M., et al., The development of vaccines against SARS corona virus in mice and SCID-PBL/hu mice. Vaccine, 2005. 23(17-18): p. 2269–72.

13. Matchett, W.E., et al., Cutting Edge: Nucleocapsid Vaccine Elicits Spike-Independent SARS-CoV-2 Protective Immunity. J Immunol, 2021. 207(2): p. 376–379.

14. Xiang, T., et al., Declining Levels of Neutralizing Antibodies Against SARS-CoV-2 in Convalescent COVID-19 Patients One Year Post Symptom Onset. Front Immunol, 2021. 12: p. 708523.

15. Li, Q., et al., Next-generation COVID-19 vaccines: Opportunities for vaccine development and challenges in tackling COVID-19. Drug Discov Ther, 2021. 15(3): p. 118–123.

16. Minor, P.D., Assaying the Potency of Influenza Vaccines. Vaccines (Basel), 2015. 3(1): p. 90–104.

17. Danaei, M., et al., Impact of Particle Size and Polydispersity Index on the Clinical Applications of Lipidic Nanocarrier Systems. Pharmaceutics, 2018. 10(2).

18. Spier, R.E., Multivalent vaccines: prospects and challenges. Folia Microbiol (Praha), 1997. 42(2): p. 105–12.

19. Yoo, K.H., et al., Possibility of exosome-based coronavirus disease 2019 vaccine (Review). Mol Med Rep, 2022. 25(1).

20. Keeton, R., et al., T cell responses to SARS-CoV-2 spike cross-recognize Omicron. Nature, 2022. 603(7901): p. 488–492.

21. Le Bert, N., et al., SARS-CoV-2-specific T cell immunity in cases of COVID-19 and SARS, and uninfected controls. Nature, 2020. 584(7821): p. 457–462.

22. O’Donnell, K.L., et al., Protection from COVID-19 with a VSV-based vaccine expressing the spike and nucleocapsid proteins. Front Immunol, 2022. 13: p. 1025500.

23. Dangi, T., et al., Combining spike- and nucleocapsid-based vaccines improves distal control of SARS-CoV-2. Cell Rep, 2021. 36(10): p. 109664.

